# The intriguing dynamics of chromatin folding and assembly

**DOI:** 10.1101/2020.02.18.953398

**Authors:** Z. Yang, W. Miao, M. R.K. Mofrad

**Affiliations:** University of Chinese Academy of Sciences; University of California, Berkeley

## Abstract

We investigate the dynamics of chromatin folding based on the “strings and binders” (SBS) model with molecular dynamics simulation. SBS model is a coarse-grained model considering a self-avoiding chain interacting with diffusive binders. By introducing transition among different categories of beads with specific transition cycles and transition probabilities, our model is capable of introducing different dynamics quantitatively during the folding process, thus capturing variety of phenomena related to chromatin dynamics. Firstly, roles of dynamics in the process of chromatin folding were examined. We discovered that there is a minimum gyration of chromatin under varying characteristic times of transition which indicates neither dramatically dynamic nor static folding process is optimal for chromatin to reach stable states with relatively low free energy. Secondly, it is noticeable that when beads transit from or into others in distinct dynamics, the equilibrium concentrations are distinct as well. As a consequence, the distribution of chromatin loop length is relevant to the dynamics of binders which can be modified by complex such as Wings apart-like protein homolog (Wapl) and SCC2/SCC4 cohesin loader complex (SCC2/SCC4). Finally, our model is able to reproduce contact matrices of both wild type HAP1 cell and ΔWAPL HAP1 cell obtained from Hi-C technology with a relatively high accuracy. Our model recapitulate the accumulating contacts at the corners of TADs and vanishing short-range contacts along the diagonal, manifesting the difference of chromatin structures before and after eliminating WAPL.

**STATEMENT OF SIGNIFICANCE:** Our model includes reciprocal transition among beads in SBS model to introduce different dynamics in chromatin folding process. Our model is able to examine the roles of dynamics in chromatin folding, reveal the loop length variation due to the concentration imbalance caused by distinct dynamics and reproduce contact matrices of both wild type and WAPL-deficient cells. Our research work provides a model to investigate the dynamics of chromatin folding quantitatively and displays its significance of revealing multiple experimental results using computational tools.

## INTRODUCTION

Three-dimensional structure of genomes inside the nucleus is a long-standing interest of biologists(1), which is essential to gene regulation(2–4). However, the mechanism behind folding and unfolding of the chromatin inside the nucleus is still under heated debate. In order to shed light on the formation of 3D chromatin structure, chromosome conformation capture (3C)-based technologies which reveal the interaction between gene loci by measuring spatial affinity in vitro have been developed in recent decades(5–8). Thanks to the rapid development of experimental technologies, the roles of multiple regulators such as cohesin, CCCTC-binding factor (CTCF), Wings apart-like protein homolog (Wapl), SCC2/SCC4 cohesin loader complex (SCC2/SCC4) in the process of assembly and disassembly of 3D chromatin architecture have been revealed (9–19). The collaboration among those regulators results in distinct dynamics of the chromatin structures(20–23).

The ring-shaped protein complex cohesin plays essential roles in the spatial organization of chromosome. Cohesin acts as a loop-extruding factor, stretching progressively larger loops to allow long-range interaction inside the chromatin. The extruding process is inclined to stalling at the boundary of topologically associated domains (TAD) due to the interaction with boundary elements. The folding process of chromatin through loop extrusion is dramatically dynamic with frequent binding and unbinding events considering both CTCF and cohesin(23). In addition, Wapl known as the antagonist of loop-extruding factors imposes an impact on the dynamic structure of chromatin by releasing cohesin from the chromatin to restrict the loop length extension(16, 21). On the other hand, SCC2/SCC4 (known as the activator) is capable of promoting the processivity of cohesin, increasing the probability of long-range interactions(16, 19). The antagonists and activators together further influence the dynamics of chromatin structure.

With the boom of experimental data such as Hi-C contact matrices, computational models based on polymer physics have been built to inversely produce the 3D structure of chromatin(24). Nicodemi et al. developed a simple polymer physics model named “strings and binders switch” (SBS) model to investigate the architectural pattern of chromatin(25, 26). Barbieri et al.(27) further utilized the model to recapitulates various experimental observation including the scaling properties of chromatin folding, the fractal state of chromatin, the processes of domain formation and looping out. One recent study(28) implemented the SBS model to explain large scale behavior of chromatin. Combined with the PRISMR algorithm developed in another work(29), the model based on polymer physics is capable of reproducing quantitative Hi-C map in a relatively high accuracy. However, these models only raised general discussion of polymer physics insights into chromatin 3D architecture and discussed little about dynamics of binders and binding sites. Here, we report a model which enables us to investigate dynamics of chromatin folding process specifically by considering quantified transition among beads of different species. The model is a straightforward extension of SBS model and the transition follows the basic rules of chemical kinetic equation. By taking the dynamic transition quantitatively into consideration, it is possible to explain the biological significance of binding and unbinding events of those key factors including CTCF and cohesin. Furthermore, we are able to prove the functions of dynamics regulator such as Wapl by comparing experimental results with our simulations.

## MATERIALS AND METHODS

### SBS model with distinct binders and binding sites

Our polymer model is based on a polymer physics model known as “String and Binders Switch”(SBS) model(25–27) in which a chromatin fiber is represented by a coarse-grained self-avoiding (SAW) polymer chain interacting with randomly sparse binders. In our model, we basically separated beads of polymer chain into 2 categories and binders into 3 categories according to interaction. More specifically, as is shown in Figure 1(a), green beads of polymer chain are sites having a larger attraction with binders named as “binding sites”, whereas blue beads within the polymer are repulsive or weakly attractive to binders named as “regular DNA”. Binding sites in our model represents DNA sequences bound with CTCF where binders reside for a longer period and regular DNA refers to DNA sequences that are not binding sites. In addition, there are three kinds of binders dynamically converting reciprocally in the model. These binders differ in the interaction with polymer chain which influence their processivity of extruding chromatin loops. The orange beads are cohesin binding with DNA chains in a regular state(without interference of activators and antagonists) which are named as regular binders (see Figure 1). Apart from regular cohesin, the cohesin activated by protein like SCC2/SCC4(16, 19) is shown as red beads in Figure 1(a) and yellow beads are representative of those cohesins released from DNA chain induced by antagonists such as Wapl. We name them as strong binders and weak binders separately.

**Figure 1:**
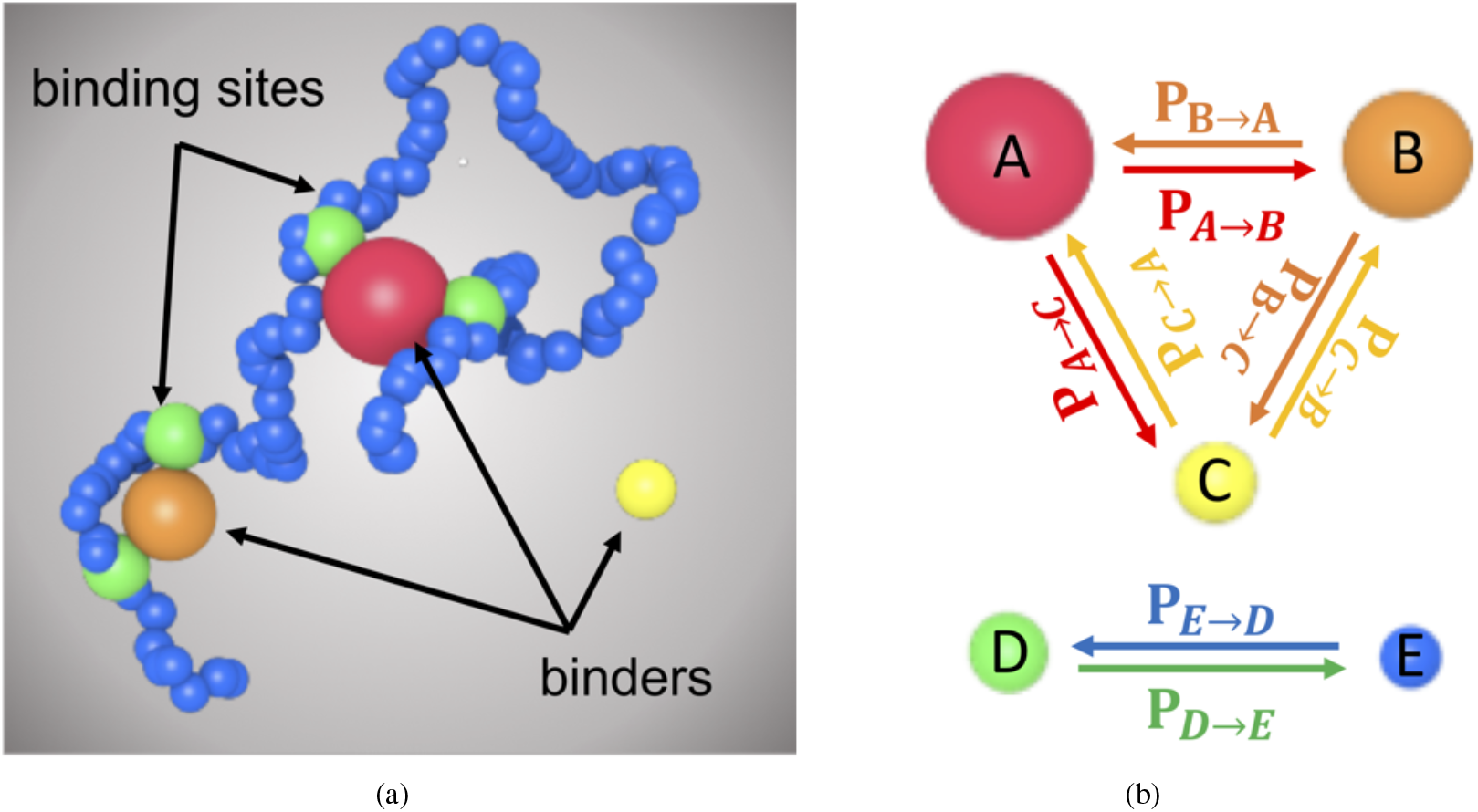
Schematic of the proposed dynamic model (a)Representation of binders and binding sites. The green beads are binding sites and the blue beads are regular DNA. The red, orange and yellow beads are strong, regular, weak binders separately. (b)Representation of transition of binders and binding sites. Transition takes place within the SAW chain or diffusive binders. The notations of transition probabilities are based on the type of beads.

As far as the interaction is concerned, we follow the original form of potential used in Chiariello's work except the parameters of interaction (*ϵ, r_c_*) between beads of strings and diffusive binders(28). Under different discussions of dynamics, we varied the interaction (*ϵ, r_c_*) and details about parameters are included under different topics in Supporting Material. The general settings used in all topics are stated below. In the model, all the beads have an identical diameter *σ*. Considering the interaction within the string, two consecutive beads are connected with a finitely extensible FENE bond(shown in Eq. 1 where *R*_0_ = 1.6*σ* and *K* = 30*K_B_T*/*σ*^2^) and the Weeks-Chandler-Andersen potential between any two beads of the string(shifted Lennard-Jones potential shown in Eq. 2 where *ϵ* = 1.0*K_B_T, r_c_* = 1.0*σ*) is included to account for excluded volume effect. In addition, the beads of the string interact with diffusive binders by the Lennard-Jones potential. From Eq. 2, it is noticeable that the interaction is determined by an energy scale *ϵ* and a cut-off distance of *r_c_*. When *r_c_* is larger than 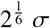, the interaction between the bead on the chain and binder is attractive, and when *r_c_* is smaller than 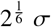, the interaction is repulsive. Moreover, a larger *r_c_* leads to a longer-range interaction. In our model, we set *r_c_* of the strong binders, regular binders and weak binders as 1. 5*σ*, 1.3*σ* and 1. 0 *σ*. As a consequence, strong binders and regular binders are attractive in different processivities whereas weak binders interact with beads of the string through repulsion. The energy scale *ϵ* for the interaction between binders and beads of the chain will be specified separately in different cases.

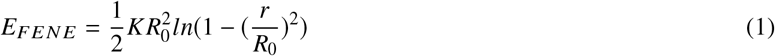

where K is the strength of the spring and *R*_0_ is distance constant representing the max length of FENE spring.

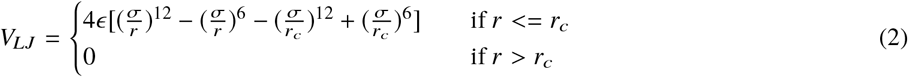

where *r_c_* is the cut off distance for the Lennard-Jones potential, *ϵ* is the energy scale of the potential.

### Dynamics quantification

To introduce different dynamics in chromatin folding, we extended SBS model within the Molecular Dynamics(MD) Simulation by allowing reciprocal transition within binders or binding sites with certain transition rate k. Transition rate k is defined by transition probability P_*i*→*j*_ and transition cycle T as 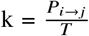. Transition probability P_*i*→*j*_ is defined as the probability of transition from category i to category j. The transition cycle is the period of transition. Two parameters are utilized to define transition rate due to the implement in MD simulations. The transition scheme and related notation are displayed in Figure 1(b). Single transition from category i to category j observes Eq. 3. The characteristic time scale of the transition is 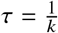, which is equal to the mean time of transition time, thus being named as period.

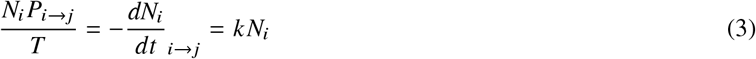

where *N_i_* is the number of category i. Here *dN_i_* only refers to the change of number from category i into category j.

In dynamic equilibrium, the concentrations of binders and binding sites are determined by the transition rates. With fixed transition cycle, the relation between concentration of beads and transition probability satisfies simple chemical kinetic equation. Eq. 4 and Eq. 5 display the relation considering binders and binding sites separately.

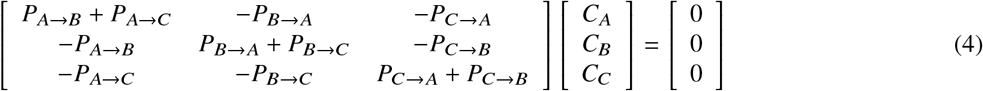

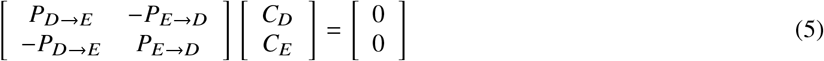

where *C_A_, C_B_, C_C_, C_D_, C_E_* are concentrations of strong binders, regular binders, weak binders, strong binding sites and weak binding sites. *P_A→B_,P_A→C_,P_B→A_,P_B→C_,P_C→A_,P_C→B_* are transition probability indicated in Figure 1(b), satisfying *P_A→B_* + *P_A→C_* < 1, *P_B→A_* + *P_B→C_* < 1, *P_C→A_* + *P_C→B_* < 1.

Apart from the transition probability, the transition period *T* plays an important role in determining the dynamics as well. A larger period of transformation leads to a less dynamic self-assembly process according to the Eq. 3. Although it is possible to obtain same transition rate *k* by varying both transition probability and transition period, we fix either transition probability or the period in order to reduce ambiguity when comparing dynamics.

## RESULTS AND DISCUSSION

### Role of dynamics

We investigated the role of dynamics in self-assembly of chromatin as dynamics is an integral part of chromatin folding, which influences various genome functions including RNA transcription and DNA replication (30). However, the general structures of chromatin under distinct dynamics are barely discussed. Figure 2(a) shows a typical example of the difference between a “static” chromatin structure (yellow line) and a “dynamic” chromatin structure (red line). More specifically, “static” refers to the situation where all transition probabilities *P_i→j_* are zero, thus characteristic time scale *τ* (defined in Materials and Methods) being infinite, whereas “dynamic” refers to the situation where *τ* is finite. As the yellow line in Figure 2(a) indicates, without dynamic transition, the chromatin falls into a metastable state (gyration around 16) quickly and is stuck all over the simulation. However, a dynamic structure of chromatin is more likely to jump out of a metastable state and fall into a state with lower free energy as the gyration shown by red line in Figure 2(a) drops more than once. The snapshot of chromatin structure in the middle is a metastable state (See Figure 2 middle panel). From the perspective of statistic mechanics, a more dynamic structure is capable of exploring more states of phase space in equal times and the probability for the system to occupy the state with a lower free energy is exponentially higher. In other words, the dynamics of self-assembly plays an important role in forming a more condensed structure of chromatin.

**Figure 2:**
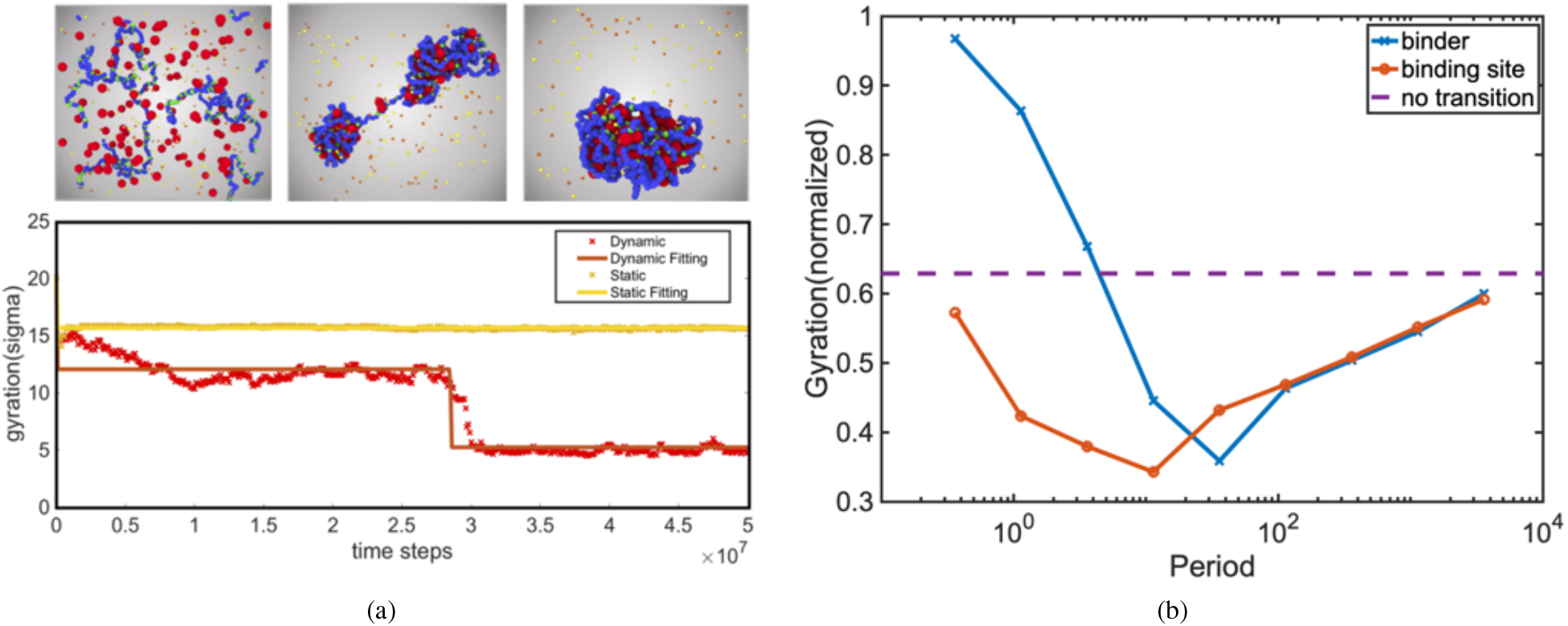
Role of dynamics(period *τ* = 0.36s, 1.11376s, 3.6s, 11.376s, 36s, 113.76s, 360s, 1137.6s,3600s) (a)Typical example of self-assembly showing the role of dynamics which is characterized by the gyration of chromatin filament. (b)Equilibrium gyration under different periods considering dynamics of binders and binding sites separately.

To further compare distinct dynamics quantitatively, we calculated the end gyrations of chromatin under different dynamics. The end gyration is an average gyration over last 2% frames of the simulation and the value is normalized according to the initial gyration. In order to vary the dynamics of folding process, we fixed the transition probability P*_i→j_* and varied the period *τ*. The specific details of dynamics setting are presented in Supporting Material. We examined dynamics of binders and binding sites separately by setting transition rate of the other category to zero (setting the category static) when considering one category. For example, there is no transition between binders when we change the dynamics of binding sites and vice versa. In both cases, as the Figure 2(b) displays, there is a minimum of end gyration as the period *τ* varies. When period *τ* is small (dynamics is dramatic), though the chromatin is able to explore more states in phase space, it is more likely to jump from a state with lower free energy to a state with higher free energy as the times of transition accumulate. When the period *τ* is large (dynamics is minor), the situation is close to a “static” self-assembly. Consequently, the end gyration is slightly smaller than that without transition. All in all, neither dramatic dynamics or minor dynamics is optimal for chromatin to fold into a condensed structure.

Despite there is a minimum end gyration for cases of both binders and binding sites, the time scales when the minimum arises are different. As the Figure 2(b) reveals, the minimum of end gyration for binders(at *τ* = 36s) arises at a larger period than that of binding sites(at *τ* = 11.376s). In addition, we noticed that when the dynamics is dramatic, the end gyration for binders is even higher than that without dynamics. In other words, the dramatic dynamics ruins the process of self-assembly for binders. However, considering the binding sites, even with the period *τ* = 0.36s, the chromatin is still more condensed than that without dynamics. These two features indicate that the transition between binding sites is more adaptable to huge dynamics than binders considering forming a close state of chromatin. The reason is that one binder (strong or regular) is always surrounded by several binding sites (DNA sequences bound with CTCF) to form many-body contact(28) given that the concentration of binding sites are higher. Even if partial binding sites are transforming into regular DNA (DNA sequences bound without CTCF), the local structure is still maintained. Therefore, transition between binding sites needs to be more dynamic to impose the same impact on the structure of chromatin as binders.

The conclusion is capable of explaining the distinct dynamics of CTCF and cohesin(23). The residence time of CTCF is ~ 1-2min while the residence time of coheisn is ~ 22min. Though given the coarse-grained nature of proposed model, any connection from MD simulation to real time is difficult to be established, the relative relation showing distinct dynamics of binders and binding sites is still conserved.

Furthermore, according to Figure 2(b), when the period is large or the dynamics is minor, the end gyrations for binders and binding sites become similar. The result manifests that in minor dynamics, dynamics itself becomes the unique factor in determining the conformation of chromatin. The reason is that when the dynamics is minor, both transitions of binders and binding sites have little impact on the structure of chromatin. As a result, instead of the category of beads that make transition, the transition frequency exerts predominant influence on the process of self-assembly.

### Loop length variation

In the last section, we investigated the impacts of overall dynamics on self-assembly of chromatin as the dynamics for different binders and binding sites were set the same. However, in general cases, the dynamics of different binders(weak, regular and strong) or different binding sites(weak and strong) are disparate. As a consequence, the concentrations of different types of beads in dynamics equilibrium are distinct according to Eq 4, thus causing variation of loop length of chromatin.More specifically, when one class of binders or binding sites is more dynamic, the concentration of the class in dynamic equilibrium is generally lower. For instance, considering ith class, when the class is more dynamic, *P_i→j_* is larger(j represents other classes except ith class). According to the Eq. 4 and Eq. 5, the concentration of ith class in equilibrium is consequently lower.

In Figure 3, we displayed the impact of distinct dynamics of binders on chromatin loop length. In order to compare different dynamics, we varied the transition probabilities of different binders while fixing the periods. Combination of larger transition probabilities for ith class of binders *D_l_* is (*P_i→j_* = 0.2, *P_i→k_* = 0.2) where *j, k* ≠ *i*. Meanwhile, combination of smaller transition probabilities *D_s_* is (P*_i→j_* = 0.1, P*_i→k_* = 0.1). In our simulations, we compared three different situations. For reference, We set *D_l_* as transition probabilities to all three classes of binders, which leads to identical concentration in equilibrium. In other two cases, we changed the combination of transition probabilities from *D_l_* to *D_s_* for strong binders and weak binders separately. Therefore, according to Eq. 4, the concentration ratios varied from 1:1:1 to 1:3:1 and 1:1:3(regular binders: strong binders: weak binders) respectively.

**Figure 3:**
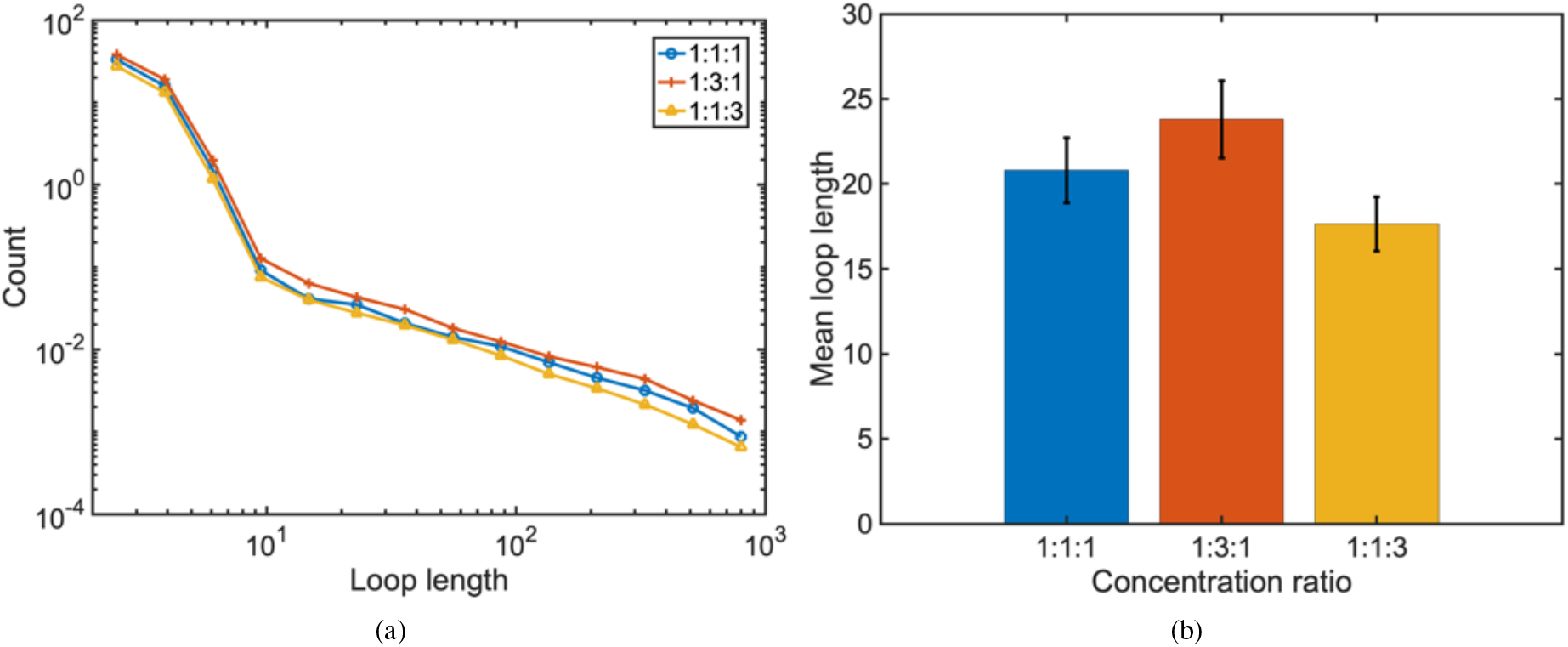
Loop length variation due to distinct dynamics of binders(concentration ratio(regular binders: strong binders, weak binders) = 1:1:1, 1:3:1, 1:1:3) (a)Distribution of loop length under different concentration ratios. (b)Mean loop length under different concentration ratios.

As a result of the distinct dynamics, we revealed the variation of distribution of loop length in Figure 3(a). The method we determined loop length is declared in Supporting Material. When the strong binders are more static, the concentration of strong binders in equilibrium increases while the concentrations of other two classes decrease, thus the overall curve of distribution of loop length being lifted comparing with the reference curve. When the weak binders are less dynamic, the distribution curve displayed an opposite variation. Figure 3(b) quantitatively provided the mean loop length under different dynamics. The results are corresponding to the distribution shown in Figure 3(a).

The conclusion that distinct dynamics are capable of varying the loop length indicates that the activity of protein related to cohesin such as Wapl and SCC2/SCC4 is able to control the structure of chromatin in nucleus. Therefore, self-assembly of chromatin is likely to be regulated by the activating or prohibiting certain proteins.

### Reproduction of Hi-C contact matrices of wild-type cell and Wapl-deficient cell

Our models were able to reproduce Hi-C contact matrices of wild-type cell and Wapl-deficient cell by varying dynamics of different binders. We selected segment(142.5Mb - 144.1Mb) of chromosome 2 of HAP1 cell whose Hi-C contact matrices has been detailed investigated (16, 31) and widely used for modeling and building new methods (32, 33).

In order to reproduce the process of folding of the chromatin segment we selected, the binding sites(CTCF sites) were assigned based on the ChIP-seq data(data accessible at NCBI GEO database(16), accession GSE95015) instead of using Hi-C map to optimize the positions of binding sites. Combined with information about TADs indicated by Hi-C contact map shown in Figure 4 (a) and (b), we split 9 categories of regular DNA and 10 types of binding sites to interact with 7 different binders(each binder interacts with one TAD). Details about the positions of binding sites and the methods we separated the beads are presented in Supporting Material. In this part, to simplify the situation, we only considered regular binders(type B) and weak binders(type C) because these two binders are sufficiently capable of representing binding(regular binders) and unbinding(weak binders) states given that Wapl is a release factor of binders (16). According to recent discoveries about the chromatin folding mechanism(10, 12, 20, 33, 34), chromatin loops are extruded to form chromosomal domains until loop extrusion process is delayed by binding sites(extrusion model). In other words, the binder is capable of attaching itself to any part of the chromatin chains during the extrusion. However, there is a larger chance that the binder is bound with binding sites when counting the contact frequency. As a result, we assigned the binders to be attractive to both binding sites and regular DNA but the attraction is stronger between binders and binding sites. In addition, we only considered the intra-TAD interaction except TAD 5. As a result, there is basically no contacts outside the triangles in Figure 4 (a). The reasons are as follows: firstly, the inter-TAD interaction is much weaker than the intra-TAD interaction; secondly, the interaction we neglected is in the regions close to the boundaries of matrices which is relatively difficult to assign because it is more likely to be influenced by segments of chromatin beyond the segment we chose. These two reasons also explained why we included TAD 5 because the interaction is relatively strong with accumulating contacts at its corner after deleting Wapl and it is considerably independent of other segments due to the fact that it is in the middle of the map.

**Figure 4:**
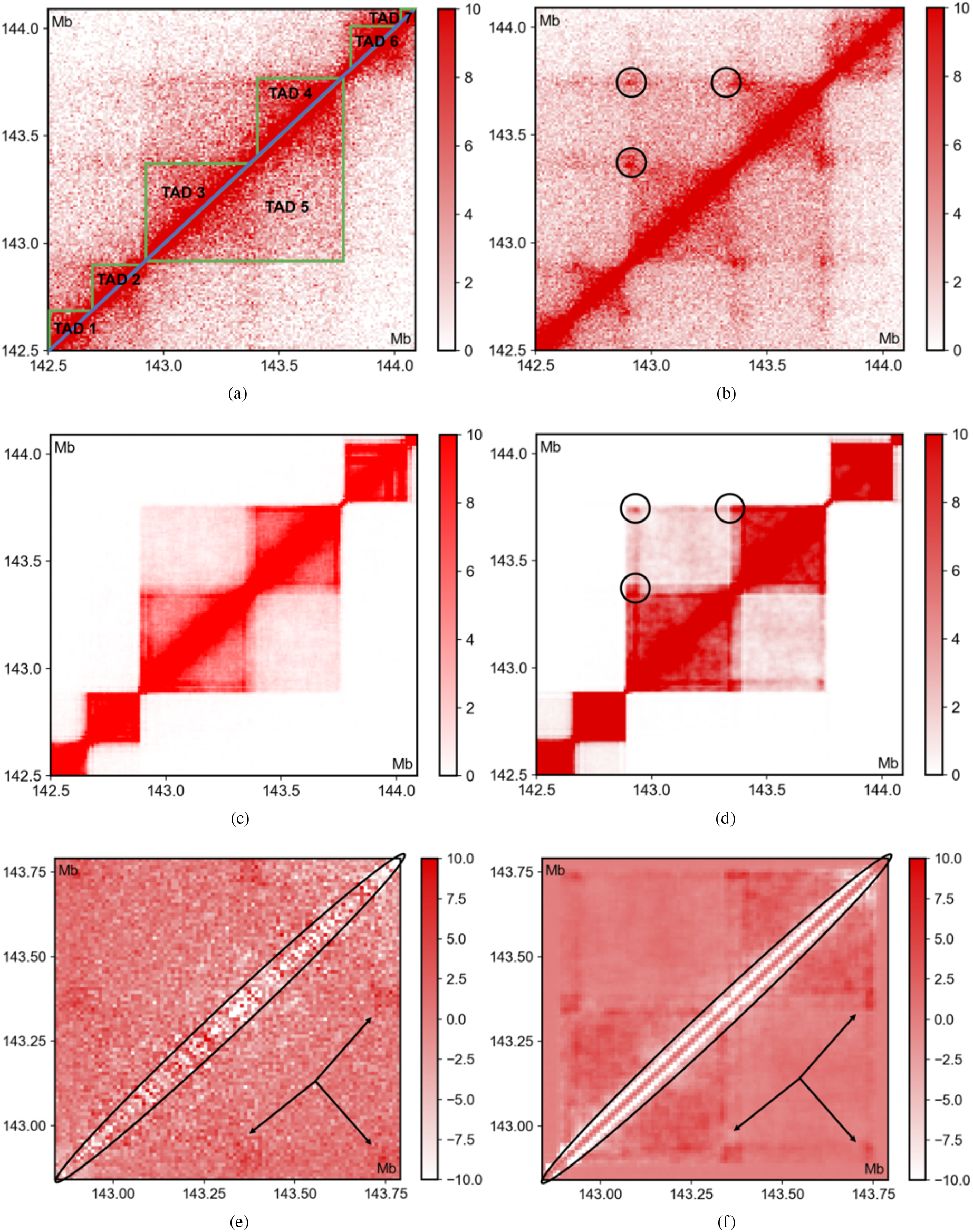
Contact matrices comparison (a), (b) Experimental Hi-C contact matrices of wild type cell and ΔWapl cell(data accessible at NCBI GEO database(16), accession GSE95015). The triangles in (a) are TADs we took into consideration. (c), (d) Simulated contact matrices for wild type cell and ΔWapl cell. Contact matrices(including (a), (b), (c), (d)) are normalized to 120000 contacts. (e), (f) Experimental and simulated matrices showing difference of contacts between two categories of cells

Figure 4 (a) and (b) revealed that erasing Wapl in cells changed the contacts distribution of chromatin by increasing the contacts at the corner of TADs and decreasing intra-TAD contact frequency(16). In order to realize the influence caused by deleting Wapl, we decreased the transition probability from regular binders to weak binders by 9 times compared with wide type cells. We assumed that the binding events were not influenced by deleting Wapl. As a consequence, the transition probability from weak binders to regular binders remain the same when simulating chromatin folding in ΔWapl cell. Details about the simulation parameters including transition probabilities are declared in Supporting Material.

According to Figure 4 (a) and (c), the model we constructed for chromatin folding of the segment of chromosome 2 of HAP1 wild type cell show great accuracy in predicting the contacts in the triangle considering we do not use any statistical methods or optimization methods for parameters. In order to demonstrate the accuracy, we used Pearson correlation r to compare the similarity of simulated contact matrix and experimental Hi-C contact matrix(29). Though we used a extremely simplified model, Pearson correlation between two maps(Figure 4 (a) and (c)) is approximately 0.92. Furthermore, through decreasing the dynamics of regular binders, we are able to simulate the contacts matrices in ΔWapl cells. As the Figure 4 (d) displays, contacts accumulate at the corner of TADs. The reason why contacts become more frequently is that when regular binders are more static, they are more likely to be stuck at binding sites due to a larger attraction. The Pearson correlation between the matrices shown in Figure 4 (b) and (d) is approximately 0.9. It is noticeable that for TAD 2 and TAD 6, the accumulating contacts at the corner are not obvious in our simulations, the main reason for that is due to that we did not include the inter-TAD interaction near the boundaries. In order to further compare the variation after deleting Wapl, we also exhibited the difference matrices of the experiment and simulation in Figure 4 (e) and (f) by subtracting maps of wide type cell from maps of Wapl-deficient cell. As the TADs near the boundaries of matrices are more difficult to predict due to the influence outside the segment we chose, we here only selected the proportion of matrices containing TAD 3, 4 and 5 to compare the difference matrices. Both matrices revealed the decrease of contacts along the diagonals(short-range interaction) and increase of contacts at the corner of TADs(long-range interaction).

However, some may argue that despite the accumulating contacts at the corner after deleting Wapl, intra-TAD contact frequency also increases while the experimental data shows an opposite result. There are actually several reasons for the difference. Firstly, Figure 4 (a) and (b) indicates that after Wapl is eliminated, the inter-TAD contact frequency increases which can cause the decrease of intra-TAD contacts owing to the normalization of total contact numbers. Nevertheless, we did not consider the inter-TAD interaction near the boundaries in our models. Secondly, we assumed that removing Wapl was not capable of influencing the binding events so the transition probability from weak binders to regular binders did not change. As a result, the concentration of regular binders in equilibrium is much too larger in ΔWapl cell than in wild type cell, leading to the increase of contacts intra TADs. Though the increase of intra-TAD contact frequency can be explained by the increase of regular binders, another question is raised: Is it also the reason for the accumulating contacts at the corner of TADs? To demonstrate that the increase of concentration of regular binders are not responsible for the increase of contacts at the corner, we further generated the difference matrices with fixed concentration of binders. The results included in Supporting Material prove that dynamics variation is the key of the accumulating contacts at the corner.

## CONCLUSION

Here we discussed about the dynamics of chromatin folding based on SBS model using MD simulations. Through varying periods and transition probabilities, dynamics of binders and binding sites were controlled and compared quantitatively in our simulations. We investigated the roles of dynamics during the process of chromatin folding and revealed that neither dramatically dynamic nor static folding is favorable for chromatin to assemble to a condensed state. In addition, we noticed that distinct dynamics is capable of causing a difference of concentration of beads involved with the folding process in equilibrium. As a result, the difference of concentration leads to chromatin loop length variation. Last but not the least, contact matrices were reproduced with relatively high accuracy based on a simply model. By decreasing the transition probabilities, difference between contact matrices of wild type cell and Wapl-deficient cell was also predicted by our simulations. All in all, our dynamic model for chromatin folding is able to reveal variety of phenomena related to dynamics of the process both qualitatively and quantitatively.

## SUPPORTING MATERIAL

Supporting Material is available at …

## AUTHOR CONTRIBUTIONS

Z.Y. and M.R.K.M. designed the research. Z.Y. carried out all simulations, analyzed the data. Z.Y.and M.R.K.M. wrote the manuscript.

