## Supporting Materials for "The intriguing dynamics of chromatin folding and assembly"

### SI Materials and Methods

### 1 Role of dynamics

In order to compare the structure of chromatin under different dynamics, we selected chains of $N_{beads} = 1000$; so for a chromosome with gene length L = 100Mb, each bead is representative of 100Kb chromatin with diameter of 87nm (1). Among 1000 beads, 20 % of them are binding sites which are randomly transiting from and into regular DNA during the simulations. As for the binders, there are 105 binders in total inside the cubic simulation cell(length = $30\sigma$). All 3 different types of binders are in the same amount. The energy scale of lj potential between binders and beads of the chain $\epsilon$ is all set to 11.67 for all three types of binders. As a result, for regular binders, the minimum of potential $E_{int} = 4.0$(1). We set the dimensional parameters of MD simulations to be the same as the former work of SBS model(1)in which viscosity $\eta = 0.1P$, time step $\Delta t = 0.012$. Considering the length scales, time unit $t = \eta(6\pi\sigma^3/\epsilon_0) = 0.036s$ under the standard physical unit (2, 3). Here, $\epsilon_0$ is the energy unit. The simulation runs for $5 \times 10^7$ time steps and the gyration average over 400 frames. The gyration is calculated over all beads within the string. Because we were investigating the impact of distinct dynamics on the process of self-assembly, especially the speed of falling into a state with relatively low free energy, we did not run extremely long simulations to assure the system reach a stable state. In addition, with a periodic transition, a longer simulation does not necessarily lead to an equilibrium state.

We discussed the dynamics of binders and binding sites separately by varying the transition probability $k$ and cycle $T$. For both binders and binding sites, transition probabilities are fixed as 10 %. As a result, cycle T is the sole variable which allows us to compare distinct degrees of dynamics without including extra factors. We selected cycle T as 0.036s, 0.11376s, 0.36s, 1.1376s, 3.6s, 11.376s, 36s, 113.76s, 360s. Correspondingly in simulation, T are 100, 316, 1000, 3160, 10000, 31600, 100000, 316000, 1000000 time steps. Consequently, the periods $\tau$ we chose are uniformly distributed in logarithmic scale.

### 2 Loop length variation

The method we calculated the distribution of chromatin loop length is as follows: firstly, two beads of the chain are regarded as forming a loop only if these two beads are spatially close and connected by the binder; secondly, we only considered the loop formed by the nearest neighbors. More specifically, if the distance between two beads is smaller than a threshold $r_t$, these two beads are spatially close. The threshold $r_t$ we set in our calculation is $2^{\frac{1}{6}}\sigma$ which is the distance at which LJ potential reaches a minimum. The threshold does change the distribution of chromatin loop length but does not influence the relative difference of distribution under different dynamics. In addition, the way we determined whether two beads of the chain are connected by binders is also checking the spatial proximity between beads of the chain and binders with same $r_t$. Moreover, the second point that we only calculated the loop formed by nearest neighbors simply assures the length of chromatin loop are repetitively counted.

As for the details of MD simulation, except the transition probabilities $k$ of different binders, other parameters were set same as the Section "Role of dynamics". We generated the distribution of chromatin loop length with over 1000 frames of simulations. Furthermore, we averaged the distribution to compare mean lengths of chromatin loop under different dynamics.

### 3 Reproduction of Hi-C contact matrices of wild-type cell and Wapl-deficient cell

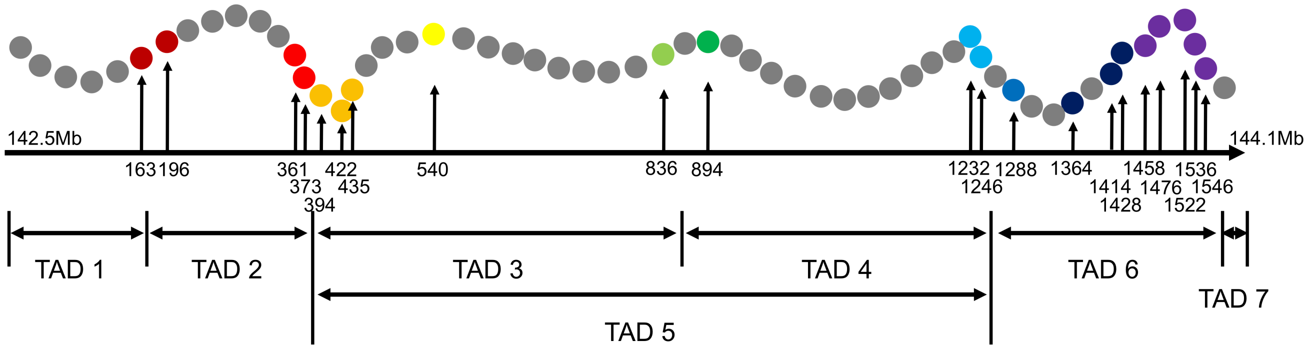


Figure S1: Arrangement of beads along the chain. Beads of different colors represent different categories of binding sites. The number in the Figure is the starting order for one binding site.

#### 3.1 Arrangement of binding sites and binders

We used chains of $N_{beads} = 1600$ to represent the segment(142.5Mb-144.1Mb) of chromosome 2. As a result, each bead is representative of 1 Kb chromatin with $\sigma = 18.75nm$. Every binding site consists of 10 coarse-grained beads as we use Hi-C map with a resolution of 10kb. The positions of binding sites are indicated by ChIP-seq data(data accessible at NCBI GEO database(4), accession GSE95015). In order to separate 7 TADs(indicated by Hi-C map), we separated binding sites into 10 different categories. For each TAD, one type of binder is assigned to interact with it and only interact with the beads inside TAD(except TAD1 and TAD7 as they are at the end of the whole chain), there are two corners which are represented by two different groups of binding sites to separate TADs. The reason we do not use same type of beads to represent two corners of TADs is due to those TADs that have overlap region(TAD3 & TAD5 or TAD4 & TAD5). According to ChIP-seq data, there are also some binding sites inside TAD but do not have such strong interaction with binders as the binding sites at the corner of TADs. We use a relatively smaller energy scale to account for that interaction. The final arrangement of binding sites is shown in Figure S1.

When it comes to the concentration of binders, it is proportional to the number of beads inside the TAD. The proportion is approximately 0.2(considering both regular binders and weak binders). For instance, if there are 100 beads inside one TAD, the number of binders interact with the TAD will be 20. However, for the TAD that is overlapped with other TADs, the number will be larger such as TAD 3 and TAD 4 in Figure S1.

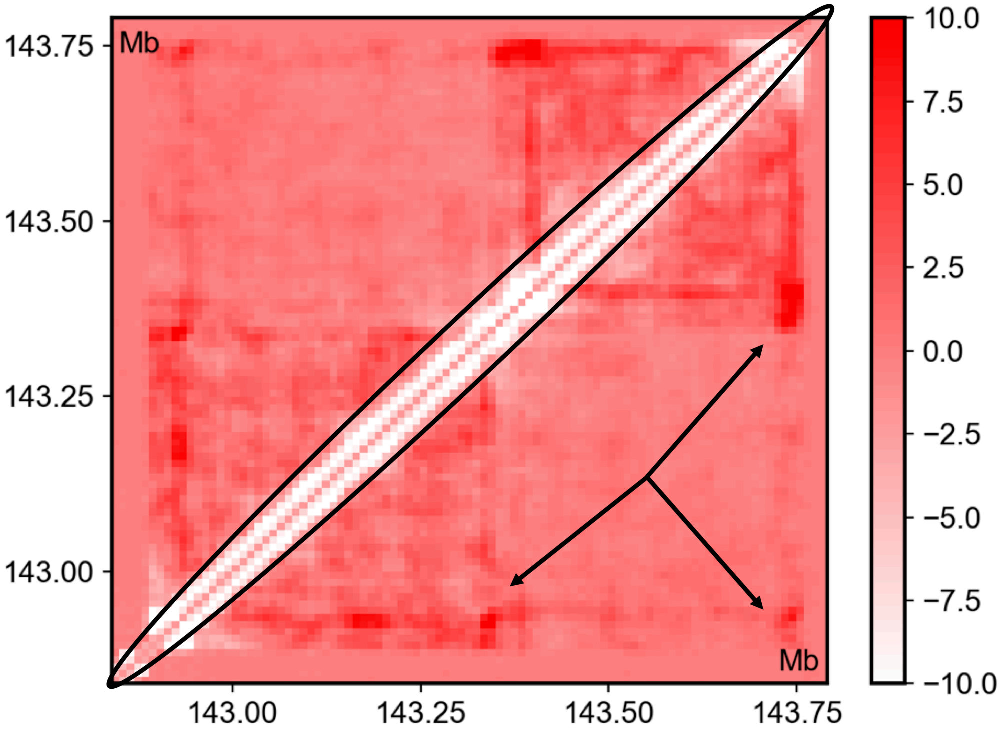


Figure S2: Simulated difference matrix without concentration variation of regular binders

#### 3.2 Parameters setting of simulations

As we have mentioned in the main body, the energy scale between binders and binding sites are stronger than the that between binders and regular DNA. So we set the energy scale between binders and DNA $\epsilon'$ to be 8.75. For other values of energy scale under 11.67, the phenomena revealed in main body are similar. In addition, we considered those binding sites as regular DNA if they are inside the TAD rather than at the corner. For instance, for TAD 5, two greens binders in Figure S1 are considered as regular DNA when interact the binders related to TAD 5. Except for these regular DNA-like binding sites for certain binders, the strength of all other interaction between binding sites and binders is set as 11.67 as before.

The parameters related to dynamics include the period and transition probabilities. We set interval for transition to be 100000 time steps. Though the volume of each coarse-grained bead is smaller than that in Section "Role of dynamics", the transition happens more frequently due to the time unit $t$ is proportional to the volume of each coarse-grained bead(relation shown in Section "Role of dynamics"). As a result, varying the volume of coarse-grained beads will not change the total volume of beads that make transition in certain time periods(real time). Furthermore, the transition probabilities were set as follows: for TAD 3,4,5, the transition probabilities $P_{B \to C} = P_{C \to B} = 0.5$ in wild type cell and $P_{B \to C} = 0.05556, P_{C \to B} = 0.5$ in Wapl-deficient cell; for other TADs, the transition probabilities $P_{B \to C} = P_{C \to B} = 0.3$ in wild type cell and $P_{B \to C} = 0.03333, P_{C \to B} = 0.3$.

#### 3.3 Concentration variation

In the main body, we showed the simulated maps with accumulated contacts at the corner of TADs under the circumstance when the different dynamics of two types of binders leads to a variation of concentration. In order to prove that the concentration variation is not responsible for the increasing corner contacts, we assumed that the deleting Wapl will also ruin the binding events. In other words, the transition from weak binders to regular binders decreases as well. To consider an extreme case, we assumed $P_{B \to C} = P_{C \to B} = 0.05, 0.03$ in Wapl-deficient cell, thus the concentration of both regular binders and weak binders remaining the same before or after eliminating Wapl. With Figure S2 indicating, increasing contact frequency at the corner of TADs and decreasing contact frequency are witnessed as well even without variation of concentration of regular binders. In conclusion, concentration variation is not responsible for the increasing corner contacts as we demonstrated in main body.
